# Deficiency in neprilysin causes laminar impairments in the visual cortex

**DOI:** 10.1101/2023.03.05.531118

**Authors:** Haneui Bae, Takao K. Hensch

## Abstract

Neprilysin (Nep) is a membrane-bound zinc-dependent endopeptidase that cleaves a wide variety of small peptides in the body. In the brain, it has gained fame as an important endogenous degrader of amyloid beta peptide, responsible for the pathophysiology of Alzheimer’s disease. Nep is expressed specifically by parvalbumin (PV)-expressing inhibitory neurons in the adult cortex, the maturation of which regulates critical period plasticity. Given that PV neuron soma and proximal dendrites are primary sites of perineuronal net (PNN) formation, we investigated the role that Nep plays in PNN and PV neuron maturation and cortical development in the mouse visual cortex, using mice whose Nep expression is constitutively knocked out (Nep KO mice). Nep expression is high in young wildtype mice (P10) and is downregulated rapidly throughout postnatal development. It is especially prominent in layer 5 where it is highly expressed by inhibitory neurons and also expressed at low levels by many excitatory neurons. Contrary to our hypothesis, Nep KO mice did not show an alteration in the relative density or gross morphology of PNNs. Instead, Nep KO mice showed reduced maximal activation of L5 after white matter stimulation and decreased number of inhibitory neurons in L4. These laminar defects may lead to impaired development of optomotor acuity in Nep KO mice.

## 2 Introduction

Neprilysin (Nep) is a membrane-bound zinc-dependent endopeptidase whose catalytic domain faces the extracellular space (**Figure 1A**). It is known by many names—neutral endopeptidase, endopeptidase-24.11, enkephalinase, membrane metalloendopeptidase (MME), CD10, and CALLA—due to its broad expression pattern in the body and its independent discoveries in various contexts (Roques et al., 1993). Nep can hydrolyze a wide variety of peptides smaller than 5 kDa such as enkephalins, substance P, atrial natriuretic peptide (ANP), and angiotensin (Matsas et al., 1984; Stephenson and Kenny, 1987; Erdös and Skidgel, 1989), but the most well-known of its substrates in the brain is amyloid beta (Aβ), the accumulation of which has been linked to the etiology of Alzheimer’s Disease (AD) (Iwata et al., 2000, 2001; Takaki et al., 2000).

**Figure 1.**
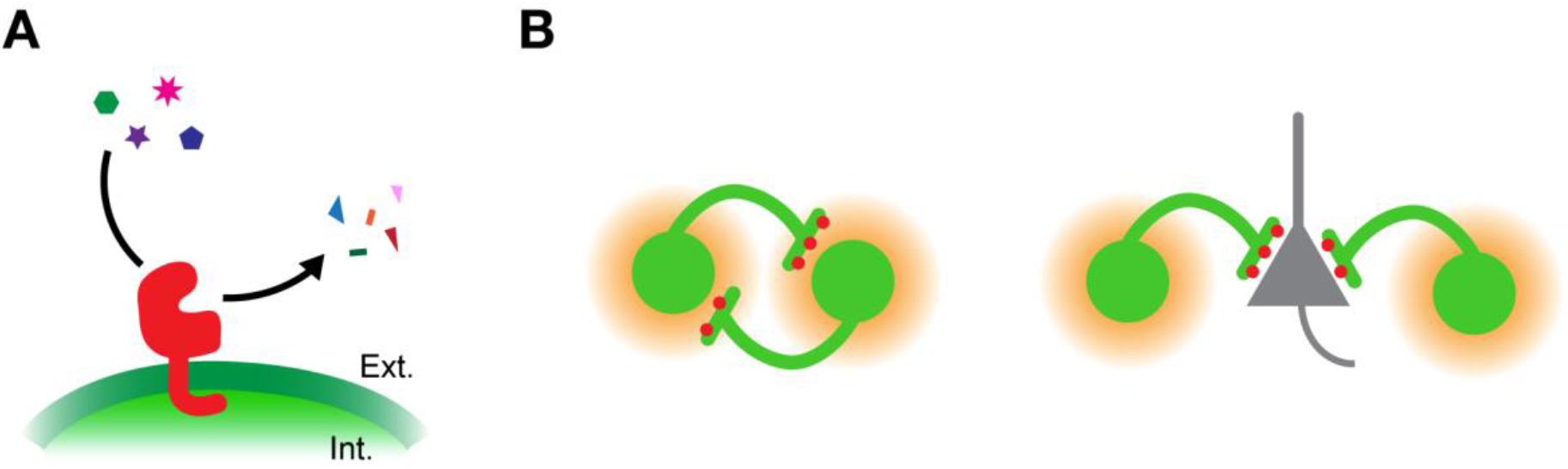
Localization of Nep in the mouse visual cortex. **(A)** A schematic depicting endopeptidase Nep (red) on PV neuron (green) plasma membrane in the mouse visual cortex. The catalytic domain of Nep faces the extracellular space, degrading a variety of small peptides. **(B)** Nep proteins subcellularly localize to the presynaptic terminals of PV neurons. The left schematic shows the potential localization of Nep to PV-PV synapses, many of which are surrounded by PNNs (orange) on PV neuron soma and proximal dendrites. The right image shows the possible localization of Nep on PV boutons onto pyramidal neurons (gray), which provide strong perisomatic inhibition.

Nep KO mice as well as WT mice infused with Nep inhibitor thiorphan showed increased levels of Aβ accumulation in the brain compared to age-matched controls (Madani et al., 2006; Mouri et al., 2006; Zou et al., 2006). In addition, Nep activity and protein levels in different regions of the brain were inversely correlated with the vulnerability to AD and decreased with age, suggesting that aging-triggered downregulation of Nep could be involved in the pathophysiology of AD (Fukami et al., 2002; Iwata et al., 2002). These findings prompted active investigations into strategies to increase Nep production and activity as a potential therapeutic to promote Aβ clearance and lower amyloid burden in the hippocampus (Iwata et al., 2004; Guan et al., 2009; Liu et al., 2009; Li et al., 2015).

The accumulation of Aβ has adverse effects not only in the aging hippocampus, but also the developing cortex. In the visual cortex, APPswe/PS1 mice, a mouse model for a familial form of AD, show impaired ocular dominance plasticity (William et al., 2012). Ocular dominance plasticity is the phenomenon in which the neurons of the primary visual cortex alter their response to the stimulation of the two eyes based on visual experience of the animal (Wiesel and Hubel, 1963). During the visual critical period of mice at postnatal day P21–35, brief 4-day monocular deprivation can dramatically reduce the cortical response to the deprived eye (Gordon and Stryker, 1996). At the peak of the critical period at P28, APPswe/PS1 mice already showed elevated Aβ compared to wildtype (WT) controls and failed to shift their ocular dominance after monocular deprivation (William et al., 2012).

Parvalbumin (PV)-expressing inhibitory neurons play an important role mediating the critical period plasticity in the cortex (Hensch, 2005). Interestingly, Nep is specifically expressed by a subtype of PV neurons in the mouse somatosensory cortex, and these neurons exhibit distinct electrophysiological properties. Nep-expressing subcluster of PV neurons are also preferentially enwrapped by perineuronal nets (PNNs) (Rossier et al., 2015), a specialized extracellular matrix associated with regulating the onset and closure of critical periods (Pizzorusso et al., 2002, 2006; Carulli et al., 2009; Beurdeley et al., 2012). Single-cell RNA-sequencing studies corroborated the finding that Nep expression is largely specific to PV-expressing neurons in the cortex, though a subset of SST-expressing neurons also express Nep (Tasic et al., 2016, 2018). This specific expression of a plasma membrane-bound ectoenzyme facing the extracellular space by PV neurons, whose soma and proximal dendrites are covered with PNNs, suggests potential interaction of Nep with the components of the net (**Figure 1B**; left image). In addition, it has been suggested that PV neuron dysfunction mediates the aberrant network activity and cognitive deficits of various AD mouse models (Verret et al., 2012; Cattaud et al., 2018; Chen et al., 2018; Hijazi et al., 2019), and that PNN-enwrapped neurons may be protected against neurodegeneration in AD (Brückner et al., 1999; Miyata et al., 2007; Morawski et al., 2012; Suttkus et al., 2012, 2014)

Given the role of Nep in the metabolism of Aβ and its specific expression in PNN-enwrapped PV neurons, the deficiency of Nep may have critical consequences in the cortical development and plasticity. We investigated this question in Nep KO mice, which harbor a whole-body deletion of the gene that encodes Nep endopeptidase (Lu et al., 1995).

## 3 Materials and Methods

### Animals

C57BL/6J mice initially obtained from Jackson Laboratories (JAX 000664) were bred in-house. Nep KO mice were obtained as a gift from Saido Lab at RIKEN BSI (first generated: (Lu et al., 1995) and were previously backcrossed to C57BL/6J to produce a uniform genetic background. Mice were maintained in heterozygous crosses, and ear-punched for genotyping at P12–21. Nep WT and KO littermates were used for experiments. Throughout the text, the term “adult mouse” refers to ages P60–150. Data from both sexes were collected and combined. Mice were housed in standard laboratory cages on a 12 hr:12 hr light:dark cycle with ad libitum access to food and water. For dark-rearing, C57BL/6J litters were housed in a light-tight box within a dark room from birth. For tissue collection of dark-reared mice, they were carried out in a light-tight container to the processing room lit with dim red light. Animal housing and experimental procedures were approved following guidelines of the Harvard University and Boston Children’s Hospital Institutional Animal Care and Use Committees.

### Amyloid Beta ELISA

Mice were anesthetized with isoflurane gas, and the brain was quickly dissected out onto cold petri dish on ice. The visual cortex was dissected using the corpus callosum anteriorly and inferior colliculi posteriorly as anatomical landmarks. To isolate the hippocampi, the brain was split down the midline through the interhemispheric fissure, and the thalamus and the underlying striatum were removed to expose the entirety of the hippocampus. The hippocampus was then carefully peeled off and cleaned of attached white matter and cortical tissue with forceps and spatulas. Dissected tissues were flash frozen in liquid nitrogen and stored at −80 °C until processing.

Frozen tissues were homogenized in 5 volumes of lysis buffer (5 M guanidine-HCl/50 mM Tris, pH8.0) by sonication and incubated on a shaker at room temperature for 4 hours. Mouse Amyloid Beta (Aβ) 40 ELISA Kit (ThermoFisher; Cat# KMB3481) was used to quantify the amount of Aβ in the tissue samples. ELISA was performed as per company protocols with the provided Aβ standards. Briefly, the standards and samples were incubated in a 96-well plate coated with antibodies against mouse Aβ40, followed by sequential incubations in solutions containing the detection antibody and secondary antibody conjugated with HRP (horse radish peroxidase). Chromogen (TMB) was added followed by the Stop solution after appropriate color development. Absorbance at 450 nm was read with a plate reader (Versamax; Molecular Devices). Relative protein amount for each sample was determined using the Micro BCA Protein Assay Kit (ThermoFisher; Cat# 23235), and the ELISA readouts were normalized to account for the differences in total protein loaded into each well. The standard curve was fit with a polynomial function, which gave a high R^2^ value (>0.99), and the values were used to convert the absorbance readings into the concentration of Aβ40.

### Quantitative Polymerase Chain Reaction (qPCR)

Mice were anesthetized with isoflurane gas, and the brain was quickly dissected out onto cold petri dish on ice. The visual cortex was dissected using the corpus callosum anteriorly and inferior colliculi posteriorly as anatomical landmarks. Dissected tissues were flash frozen in liquid nitrogen and stored at −70 °C until processing.

Frozen tissues were submerged in TRIzol™ reagent (Thermo Fisher Scientific) and mechanically homogenized with a pellet pestle (Cordless pestle motor and sterile disposable pestles, VWR International). After addition of chloroform and vigorous mixing, samples were centrifuged for 15 min at 12000 × *g* at 4 °C. Clear supernatant was carefully transferred to RNeasy Mini columns (Qiagen), and the rest of RNA extraction was carried as per supplier instructions. RNA was eluted in nuclease-free water, quantified using NanoDrop 1000 Spectrophotometer (Thermo Fisher Scientific), and stored at −70 °C until use. RNA was reverse transcribed using High-Capacity RNA-to-cDNA™ kit (Thermo Fisher Scientific). qPCR reactions were set up using TaqMan™ Gene Expression Master Mix on MicroAmp™ Fast Optical 96-Well Reaction Plates (Thermo Fisher Scientific) and run on StepOnePlus Real-Time PCR System (Applied Biosystems). TaqMan probe used: Mm00485028_m1 (*mme*; Neprilysin). All results were normalized to GAPDH expression and then to WT level per gene.

### In Situ Hybridization

#### Tissue collection and sectioning

The dissection bench was cleaned with RNase AWAY™ Surface Decontaminant (ThermoFisher Scientific) to minimize contamination with RNases. Mice were anesthetized with isoflurane gas, and the brain was quickly dissected out onto cold petri dish on ice. After a few washes with cold sterile saline, the brain was submerged into a mold filled with Tissue-Tek^®^ O.C.T. Compound and placed on dry ice until frozen. The fresh-frozen brain blocks were stored at −70 °C until cutting.

Brain blocks were equilibrated inside the cryostat (Leica CM1950) set at the following temperatures, chamber −14 °C and specimen block −12 °C, for at least one hour before sectioning. 25-μm sections encompassing the primary visual cortex were cut, captured on Superfrost^®^ Plus Micro Slides (Cat# 48311-703; VWR^®^), and immediately frozen on dry ice. The sections were stored at −70 °C until processing.

#### In situ hybridization (RNAscope^®^)

mRNA transcripts were labeled and visualized with RNAscope^®^ Fluorescent Multiplex Assay (ACD Bio) as per company protocols. Briefly, frozen sections were fixed in 4% paraformaldehyde (PFA) and dehydrated in increasing concentrations of ethanol. After pretreatment, slides were incubated at 40 °C with probes targeting the transcripts of interest. The following probes were used for P40 sections—Mm-Mme (Cat# 415421), Mm-Pvalb-C2 (Cat# 421931-C2), Mm-Gad1-C3 (Cat# 400951-C3), Mm-Gad2-C3 (Cat# 439371-C3)—and P10—Mm-Mme, Mm-Slc17a7-C2 (Cat# 416631-C2), Mm-Gad1-C3, Mm-Gad2-C3. The sections were then treated with a series of amplification reagents, stained with DAPI, and coverslipped with mounting medium (ProLong™ Gold Antifade Mountant). The slides were sealed with nail polish and stored in dark at 4 °C as soon as the nail polish was dry until imaging.

### Immunohistochemistry

Mice were deeply anesthetized with an overdose of pentobarbital (5 mg/20 g; Euthasol^®^, Virbac) and perfused intracardially with cold saline followed by ∼40 mL 4% PFA in 0.01 M phosphate buffered saline (PBS) (freshly diluted from 32% EM Grade PFA; Electron Microscopy Science). The brains were dissected out and post-fixed in 4% PFA at 4 °C for 24 hr. The duration of post-fix was kept as accurately as possible (±30 min) for consistencies in immunostaining with antibodies sensitive to the extent of fixation. After post-fix, the brains were washed 3 times and stored in PBS at 4 °C until sectioning (usually within 1–3 days). Serial coronal sections of 40-μm thickness encompassing the visual cortex were cut on a vibratome (Leica VT1000S) and stored in 96-well flat-bottom plates filled with 0.02% sodium azide PBS at 4 °C (Santa Cruz Biotechnology).

For each round of immunohistochemistry experiments 2–3 cortical sections (4–6 hemispheres) were used for each animal to account for within-animal variability. Coronal sections containing binocular primary visual cortex were chosen based on the distance from the section in which the white matter of each hemisphere connects in the corpus collosum. The sections were incubated in BSA block solution (bovine serum albumin; 5% BSA, 0.5% Triton X-100, 0.01M PBS) for 1–2 hr at room temperature (RT), before 24–48-hr primary antibody incubations on a shaker at 4 °C. Primary antibodies used were: rabbit-α-PV (1:500; Swant), biotinylated Wisteria Floribunda Agglutinin (bWFA) (1:500; B-1355, Vector Laboratories), rabbit-α-somatostatin14 (1:500; T-4103, Peninsula Laboratories International, Inc.), and guinea pig-α-GABA (1:100; AB175, Millipore Sigma). The sections were washed 3 times for at least 10 min each with 0.1% Triton X-100 in PBS (PBST) before overnight incubations in secondary antibodies at 4 °C. Secondary antibodies used were: goat-rabbit-AlexaFluor488, goat-mouse-AlexaFluor594, streptavidin-647, and DAPI (all at 1:1000 dilution; Thermo Fisher Scientific). Primary and secondary antibody dilutions were made in BSA block solution. After washing with PBST and PBS, the sections were mounted on slides with Flouromount-G^®^ mounting medium (Cat# 0100-01, SouthernBiotech). Slides were allowed to dry protected from light overnight at RT, sealed with nail polish, and transferred to 4 °C for storage.

### Image acquisition and analysis

#### Acquisition

Images were acquired using ZEISS LSM 880 Confocal Microscope at Harvard Center for Biological Imaging. Tiled images were centered around the binocular visual cortex, whose boundary between secondary visual areas was identified by the thickening of the cortex especially layer 4 (L4) and the presence of intense WFA staining. All cortical layers were imaged. The parameters of image acquisition, such as laser power and gain, were determined on the first image and kept the same for all slides in the experiment. When possible, all images were acquired in one imaging session on the same day, and at maximum over 3–4 days for consistency.

#### Analysis

Tiled images were stitched with Zen software (ZEISS) and converted to TIFF files using Fiji (Schindelin et al., 2012). To blind the experimenter to the genotype and identity of each image, the files were randomly renamed with numbers using a custom Python code. At the end of all quantifications, each image was re-matched with its identity for statistical analysis.

For immunohistochemistry images, a cortical region 450-μm wide in the binocular visual cortex was used for analysis. Within this region, cortical layers were delineated using DAPI signals. L4 was identified by a band of strong DAPI staining in the middle of the cortex due to the increased density of cells. L6 is characterized by a slight increase in cell density compared to L5 above. L1 is almost devoid of cell bodies. A region of interest (ROI) for each layer was drawn in Fiji.

The number of immunopositive cell bodies were counted using Fiji Cell Counter plug-in. The number of double-positive cells were counted to calculate the fraction of overlap. For each animal, 4– 6 sections were stained and imaged. The counts for each image were added together to give one value per animal, and the fractions were averaged. Raw cell counts were normalized by the area of summed layer ROIs and shown as density. Quantified values across the five cortical layers between WT and KO were compared with Repeated Measures Two-way ANOVA and post-hoc test with Bonferroni correction.

For *in situ* hybridization images, punctate signals of *Mme* were first thresholded and masked to remove the background signal. To count the number of *Mme*-positive cells, a cluster of *Mme* puncta that fall within the boundaries of *Gad* or *Slc17a7* were counted as one positive cell. There were almost no clusters that fell outside of the boundaries of *Gad* or *Slc17a7*. Cell were counted, normalized, and compared as above.

### Voltage-sensitive dye imaging (VSDI) and analysis

#### Solutions

Two types of artificial cerebrospinal fluid (ACSF) solutions were used: one for recording and one with low calcium and high magnesium for cutting and dye incubations. Cutting ACSF contained (in mM): 130 NaCl, 24 NaHCO_3_, 10 Glucose, 3.5 KCl, 1.25 NaH_2_PO_4_, 5 MgCl_2_, and 1 CaCl_2_. Recording ACSF was the same except for 1.5 MgCl_2_, and 2.5 CaCl_2_. 10x standard ACSF solution was kept at 4°C, and MgCl_2_, and CaCl_2_ were freshly added on the day of the experiment. The voltage-sensitive dye Di-4-ANEPPS (Cat# D1199; Thermo Fisher Scientific) was dissolved in dimethyl sulfoxide (DMSO) to a stock concentration of 5 mg/mL and was diluted to a final concentration of 5–10 μg/mL in cutting ACSF. All solutions were continuously bubbled with carbogen (95% O_2_, 5% CO_2_).

#### Dissection and Recording

Mice were anesthetized with isoflurane before decapitation. The brain was quickly taken out of the skull and fully submerged in ice-cold cutting ACSF during the dissection. A cut was carefully made in the frontal cortex to create a base to glue the brain to the vibratome platform. In order to preserve the cortical circuits perpendicular to the pial surface of the visual cortex, the cut was angled slightly to match the cortical curvature in the occipital lobe. 300-μm-thick slices were cut on the vibratome (Leica VT1000S) at frequency 8 (80 Hz) and speed 2 (0.075 mm/s). Appropriate visual cortical slices were identified by the characteristic shape of the white matter. Slices were recovered in 33 °C cutting ACSF solution for 30 min and kept at RT until recording.

Slices were sequentially transferred to the VSD solution to ensure consistent level of dye absorption and incubated for 90–150 min at RT until they reached a background fluorescence intensity of 500– 900 (artificial units used in MiCAM ULTIMA software). The dye absorption of slices depended both on the concentration of the dye and the duration of incubation.

The slices were then transferred to recording chamber filled with recording ACSF (20–22 °C) and imaged using an Olympus MVX10 microscope with a 1x 0.25 NA objective. Stimulation (50–500 μA, 1 ms pulse) of white matter with an ACSF-filled glass pipette was controlled by a programmable pulse generator (MiCAM ULTIMA; SciMedia) linked to a constant current stimulus isolation unit (Iso-Flex; A.M.P.I.). Excitation light from a shuttered 150-W halogen lamp (MHF-G150LR; Moritex) was band-pass filtered (515–535 nm) and reflected toward the sample by a 570-nm dichroic mirror. Emitted fluorescence, reflecting a change in potential across membranes (Grinvald and Hildesheim, 2004), was long-pass filtered (590 nm) and imaged using a MiCAM ULTIMA CMOS-based camera (SciMedia). Changes in fluorescence (compared to the background fluorescence at 0 ms; DEF acquisition mode) over 511 ms were recorded at 1000 Hz and averaged across 10 trials.

After imaging, cortical slices were fixed in 4% PFA at RT for 30–50 min, washed, and stored in PBS at 4 °C. They were stained with DAPI for 1 hr in RT as described above. To mount and coverslip 300-μm sections without damaging the tissue, a reservoir was created with two layers of parafilm (∼260-μm thick) around the tissue. Slides were imaged with ZEISS Axio Scan.Z1 Slide Scanner (Harvard Center for Biological Imaging).

#### Analysis

Analysis of the VSDI data was performed on MATLAB (MathWorks). Data were converted from MiCAM ULTIMA output files (.rsd, .rsh, and .rsm files) to a MATLAB-readable format using Rhythm, a program developed by Efimov Lab at Washington University in St. Louis (Matt Sulkin, Jake Laughner, Xinyuan Sophia Cui, and Jed Jackoway; obtained from GitHub). Each data file consisted of a 100x100 pixel background image at 0 ms, and a 100x100x511 array containing the fluorescence measurement at every ms for 511 ms.

The data array was first smoothed by convolution with 3x3x3 box kernel (smooth3). For each pixel, baseline was calculated by averaging the values at 50–90 ms (stimulus onset at 100 ms). Fluorescence change at each pixel was then normalized to the baseline fluorescence to calculate dF/F. To more easily visualize and analyze activity in each cortical layer, images were flattened by moving background pixels from the top of the image to the bottom for each column. Background pixels were defined as any pixels in the background image with fluorescence values lower than one standard deviation from the mean of all background values. A peak value for each pixel was determined by calculating the maximum value that occurs during 100–120 ms. Any “peak” values less than 2 standard deviation from baseline values were considered noise.

To overlay laminar information from DAPI-stained images to VSDI data, the depth of each layer boundary was divided from the total depth of the cortex to get a fraction depth and was multiplied to the total number of pixels for the cortex in VSDI images (39–40 pixels). The layers delineated in this manner matched well with the VSDI activity patterns. For a typical slice, L1 ranged from pixels 1 to 4, L23 5–14, L4 6–20, L5 21–31, and L6 32–40.

To analyze layer current/response curves, a region of interest (ROI) for each layer was chosen from the center of vertical spread of activity. Each ROI was 3 pixels wide. For L23, L5, and L6, ROIs were 5 pixels high, selected from the middle of the layer and avoiding the edges. L4 ROIs were 3 pixels high. Peak dF/F of all pixels contained in the ROI was averaged and graphed for different current stimulation intensities. To account for small differences in staining intensity and the actual current output from the tip of the glass pipette in different experiments, the dF/F values of L4, L5, and L6 are normalized to the maximum value in L23 for each slice, while the values of L23 are shown without normalization.

These curves took a sigmoidal shape, with responses that increased linearly in the middle and eventually saturated at high current stimulations. These current/response curve for each slice were fit with a sigmoidal curve (using GraphPad Prism, outlier removal coefficient Q = 1%), with four parameters, Top, Bottom, EC50, and HillSlope. Bottom was constrained to a value of 0 because the baseline had already been subtracted from each data point, and HillSlope was constrained to a value between 0 and 0.4 to aid convergence. HillSlope taken by most fits was around 0.01 and did not differ between WT and KO. The remaining two parameters, Top, the upper asymptote that represents the maximal response of the slice, and EC50, the current at which half maximal response occurs, were used for comparison between WT and KO mice (t-test). Upper/lower ratio was calculated by dividing the L23 value by L5 value (unnormalized) for each slice at each current.

For horizontal spread analysis, dF/F values at each X position in L23 and L4 were averaged and cropped so that the center of the vertical column of activity was at 0. Half maximal value of L23 was set as the threshold, and the width of spread for both L23 and L4 curves was defined as the number of continuous pixels greater than the threshold. The widths of L23 and L4 were then compared between WT and KO (t-test).

### Optomotor task for visual acuity

This experiment was performed by Georgia Gunner of Neurodevelopmental Behavior Core, Boston Children’s Hospital, as described in (Prusky et al., 2004). Optomotor testing apparatus consisted of a raised circular platform (13 cm from the floor) in a box (39 × 39 × 32.5 cm) surrounded by four computer monitors that displayed the visual stimulus. The floor and the ceiling of the box were covered with mirrors, which created an illusion of an infinite cliff. A camera was positioned directly above the platform to monitor the head movements of the mouse. A computer program (OptoMotry; CerebralMechanics, Lethbride, Alberta, Canada) projected on the monitors sine wave gratings with various spatial frequencies (0.03–0.5 cycles/degree; 100% contrast) slowing moving at 12 degrees/s. Visual stimuli were extended by the floor and ceiling mirrors, and from the perspective of the platform, the monitors appeared as windows overlooking a surrounding 3-D world.

Mice were placed one at a time on the platform and allowed to move freely. When a moving grating perceptible to the mouse was projected on the wall, it normally stopped moving and began to track the grating with reflexive head movements in concert with the rotation. An experimenter assessed whether the animals tracked the cylinder by monitoring the video. All animals were habituated before testing by handling and placing them on the platform for a few minutes at a time. To minimize handling stress in young mouse pups, developmental optomotor acuity was measured on non-consecutive days from multiple litters. The number of animals used per age is shown in **Supplementary Table S1**.

### Statistical analysis

All data are expressed as mean ± SEM. Statistical tests were performed using GraphPad Prism (Repeated Measures Two-Way ANOVA, Two-Way ANOVA, t-test, and sigmoidal curve-fitting). Post-hoc analyses were performed with Bonferroni correction. For comparisons that were significant, the test used and the significance levels (* *p* < 0.05, ** *p* < 0.01, *** *p* < 0.001, **** *p* < 0.0001) are reported in the figures.

## 4 Results

### Nep expression becomes downregulated and PV neuron-specific over development

Protein Nep is synthesized by the *Mme* (membrane metallo-endopeptidase) gene in mice. To understand the developmental expression level of *Mme* in the mouse visual cortex, we first measured the level of *Mme* expression in homogenized visual cortical tissue of C57BL/6J mice. *Mme* expression was the highest at postnatal day 10 (P10) and decreased with age after eye-opening around P14 (**Figure 2A**; solid line). Interestingly, this developmental decrease of *Mme* expression was experience-dependent, as rearing mice in complete darkness from birth prevented the downregulation and the animals maintained a P14 level of expression (**Figure 2A**; dashed line). Consistent with this expression pattern, Nep KO mice show an elevated level of Aβ40 protein, one of the major substrates of Nep in the brain, both in juvenile and adult visual cortex (**Figure 2B**).

**Figure 2.**
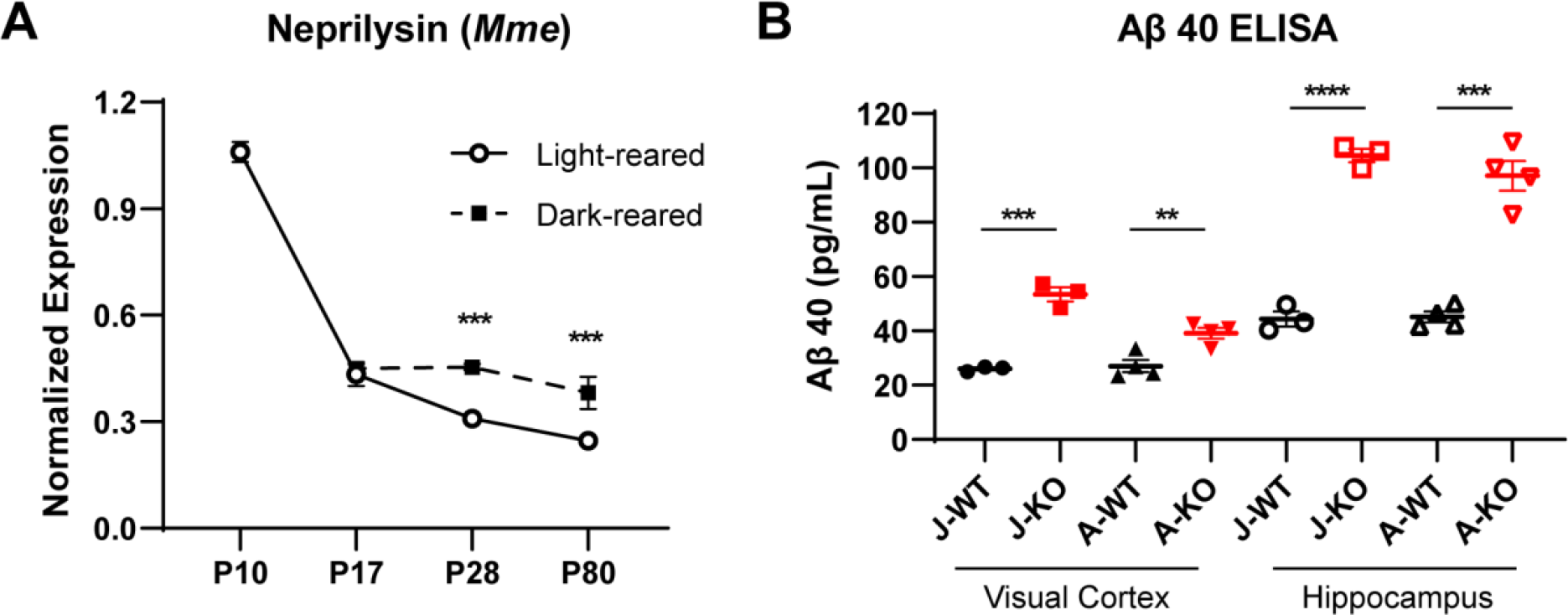
Developmental expression of Nep and Aβ40. **(A)** Developmental expression of Nep (*Mme*) in the mouse visual cortex. In the visual cortex homogenate, *Mme* expression is the highest at P10 and showed an age-dependent decrease of expression. Dark-reared mice maintained a higher expression level compared to light-reared animals. Results are expressed as mean ± SEM. (Light-reared n = 4–6, dark-reared n = 3–5; Two-way ANOVA with Bonferroni post-hoc test, *** *p* < 0.001) **(B)** Nep KO mice (red) showed a higher level of Aβ40 protein in the visual cortex, as well as the hippocampus, compared to WT (black). Both juvenile (J; P34) and adult (A; P125–130) KO mice showed the elevation in Aβ40. (Juvenile WT n = 3, KO n = 3; Adult WT n = 4, KO n = 4; t-test, ** *p* < 0.01, *** *p* < 0.001, **** *p* < 0.0001)

The elevated expression of *Mme* in P10 mouse pups may indicate the following: a higher level of expression in PV neurons with the same pattern of expression as in adult mice (Rossier et al., 2015), nonspecific expression of *Mme* by different cell types, or a combination of the two. To differentiate between these possibilities, we performed *in situ* hybridization for *Mme* transcripts, as well as for cell type-specific markers *Pvalb*, *Gad*, and *Slc17a7* at P10 and P40 (**Figure 3**). *Gad* produces the synthesis enzyme for the inhibitory neurotransmitter GABA and is a marker of inhibitory neurons, and *Slc17a7* encodes the transporter of the excitatory neurotransmitter glutamate, Vglut1. *Pvalb* expression in PV neurons only begins after eye-opening and is absent at P10 (Río et al., 1994). At P40 visual cortex, *Mme*+ puncta were found concentrated in *Pvalb*-expressing cells distributed evenly throughout cortical layers L23–L6 (**Figure 3C**; top, blue arrows), with minimal expression in L1 (**Figure 3A**). At P10, *Mme* expression was highly elevated in all layers of the cortex, but most prominently in L5 (**Figure 3B**; P40 158.02 ± 11.83 cells per mm^2^ vs. P10 642.49 ± 35.22). P10 L5 visual cortex showed *Gad*-expressing cells bodies with very high concentration of *Mme* transcript (**Figure 3C**; bottom, blue arrows) among a more diffuse distribution of *Mme* in *Slc17a7*-expressing cells (blue arrowheads). The fraction of *Gad* and *Mme* double-positive cells among all *Mme*+ cells is only 0.340 ± 0.009 (**Figure 3D**; dashed line). This is in contrast to P40 L5, where the majority of *Mme* puncta were contained within cells expressing *Gad* (0.869 ± 0.024; solid line) and *Pvalb* (0.747 ± 0.015). Also in L6, the fraction of *Gad*/*Mme* colocalization was low in P10 (0.371 ± 0.032) compared to P40 (0.973 ± 0.016). The drop is because a large number of *Slc17a7*+ cells near the border of white matter lowly express *Mme* (**Figure 3A**).

**Figure 3.**
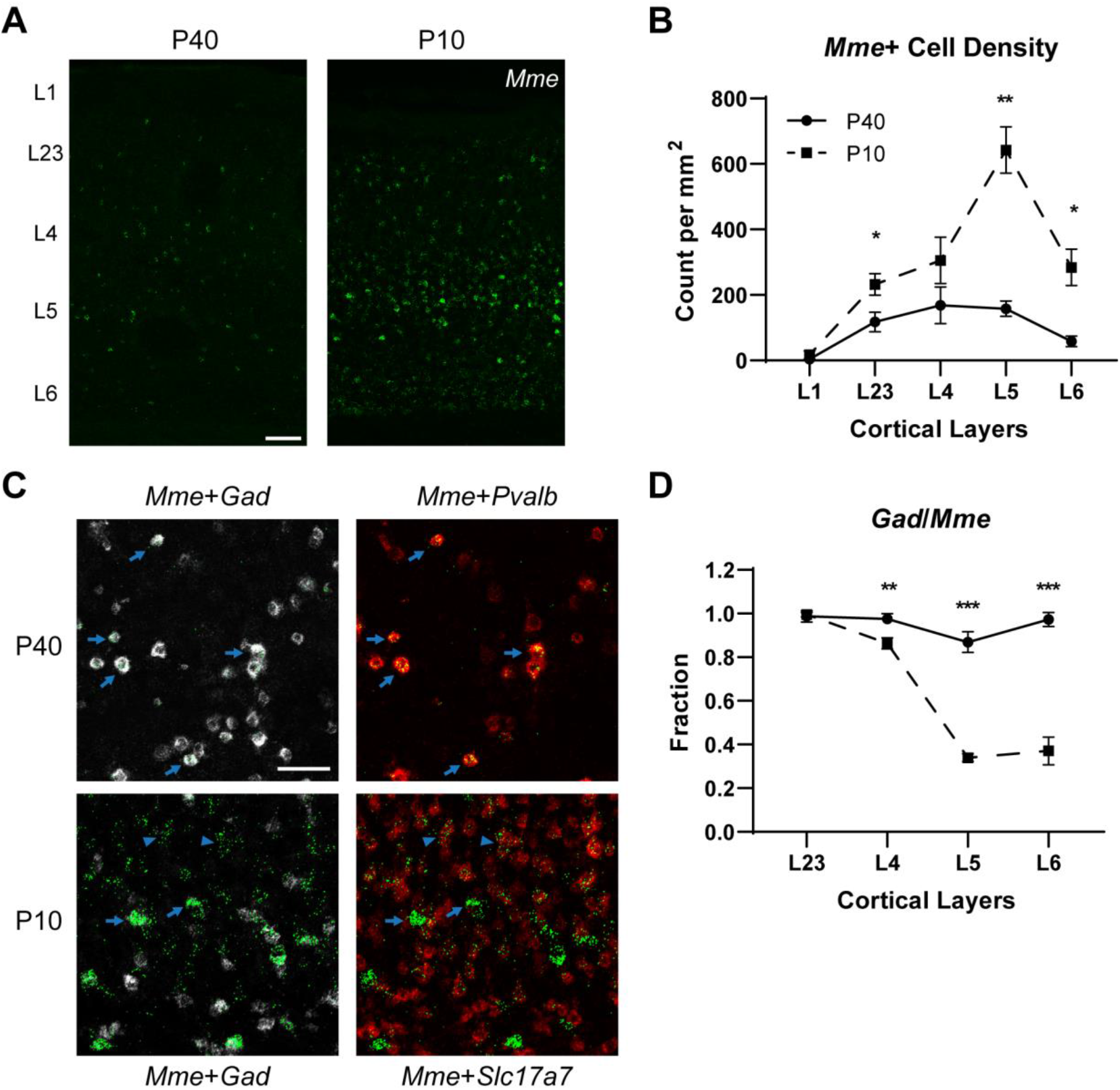
Laminar and cell type-specific expression of *Mme* in the mouse visual cortex. **(A)** Representative images showing *Mme in situ* hybridization signals in P40 and P10 C57BL6/J visual cortical sections. Cortical layers (L1–L6), determined by DAPI staining (not shown), are delineated by white dashed lines. (Scale bar, 100 μm) **(B)** Quantification of *Mme*+ cell density throughout cortical layers in P40 and P10 visual cortex. Larger number of cells express *Mme* at P10 compared to P40 especially in L5 (P40, 158.02 ± 11.83 vs. P10, 642.49 ± 35.22) of the visual cortex. (P40 n = 4, P10 n = 4; Repeated Measures Two-way ANOVA with Bonferroni post-hoc test, * *p* < 0.05, ** *p* < 0.01) **(C)** Higher magnification images showing L5 of P40 and P10 visual cortex. While *Mme*+ cells (green) are mostly *Gad*-expressing inhibitory neurons (gray) in P40 (the majority of which are *Pvalb*-expressing; red, top; blue arrows), only a fraction of them are inhibitory at P10 (blue arrows). Most of the non-inhibitory *Mme*+ cells are *Slc17a7*-expressing excitatory neurons (red, bottom; blue arrowheads). (Scale bar, 50 μm) **(D)** Quantification of the fraction of *Mme*+ cells that express *Gad* throughout cortical layers in P40 and P10 mouse visual cortex. (P40 n = 4, P10 n = 4; Repeated Measures Two-way ANOVA with Bonferroni post-hoc test, ** *p* < 0.01, *** *p* < 0.001)

These data show dramatic changes in the level and pattern of Nep expression over early postnatal development. There is an overall reduction in the level of Nep expression in the visual cortex, and this change is much more prominent in the lower layers. Different cell types also portray distinct expression patterns; while the primary producers of Nep are inhibitory neurons in both P40 and P10 visual cortex, excitatory neurons in the lower layers of the cortex also show diffuse expression at P10. This more global expression pattern diminishes over development and becomes largely absent by P25 (**Supplementary Figure S1**), resulting in the adult distribution consistent with previous reports (Rossier et al., 2015).

### The density of inhibitory neurons is reduced in L4 of Nep KO mice

The early high expression and later specific expression of Nep in PV neurons suggest potential roles of Nep in the postnatal development of the cortex. We investigated this in mice where Nep is globally knocked out (Nep KO) (Lu et al., 1995). PV and PNNs are markers of cortical maturation in postnatal development. PV and WFA immunostaining of adult Nep WT and KO mice showed a 17% decrease in the number of both PV and WFA cells in layer 4 (L4), but not in other cortical layers (**Figure 4**). The fraction of WFA-enwrapped PV cells and WFA-positive cells that were PV neurons was not different between Nep WT and KO (**Figure 4B**), which suggested that the reduction in the number of WFA cells was driven by the drop in PV cells. Gross morphology of the nets and the intensity of WFA staining in L4 were not altered (WT 83.56 ± 2.81, KO 77.90 ± 0.76, n=4; t-test, *p* = 0.176).

**Figure 4.**
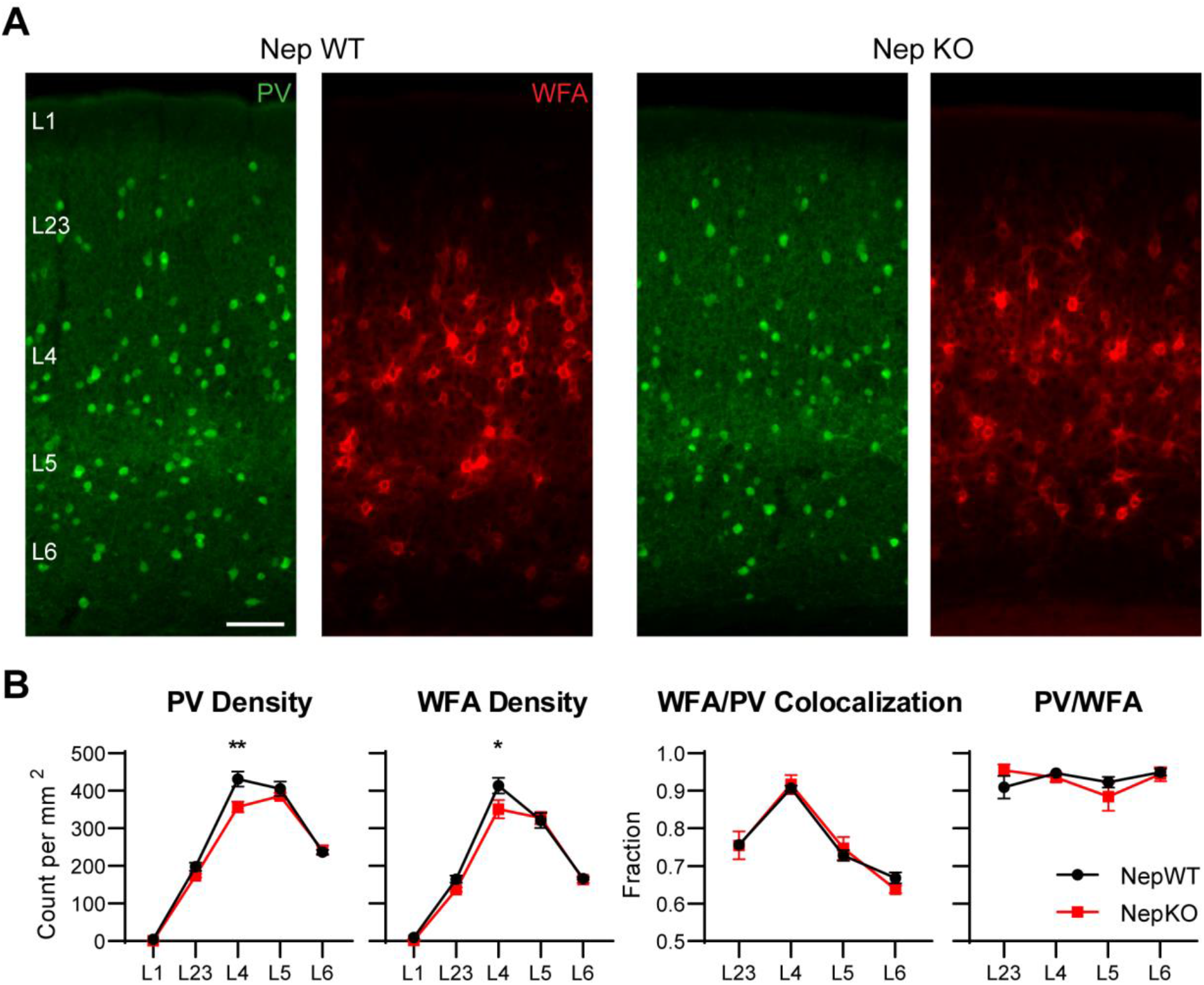
Decreased numbers of PV and WFA cells in L4 of adult Nep KO mice. **(A)** Representative images showing L1–L6 of V1 in Nep WT and Nep KO. (Scale bar, 100 μm) **(B)** Quantification of PV+ and WFA+ cells in each layer of the cortex. Adult Nep KO mice show decreased PV and WFA cells specifically in layer 4 of binocular visual cortex. The proportion of PV neurons that are enwrapped by PNNs is unchanged in the KO mouse. (Nep WT n = 7, KO n = 4; Repeated Measures Two-way ANOVA with Bonferroni post-hoc test, * *p* < 0.05, ** *p* < 0.01)

The drop of PV cell density in L4 could be a PV-specific defect or a result of a general reduction in the number of inhibitory cells in L4. Inhibitory neurons can be divided into three mostly nonoverlapping subpopulations expressing PV, SST, and serotonin 3A receptor (5-HT_3A_R) (Rudy et al., 2011). Among these 5-HT_3A_R-expressing interneurons primarily localize to L1 and L23 of the cortex, so inhibitory neurons in L4 are expected to be mostly PV and SST cells. Immunostaining for SST and GABA showed that there is a similar 17% drop in the number of SST+ and GABA+ cells specifically in L4, revealing that this is a general inhibitory cell defect (**Figure 5**). The phenotype was also present in P17 and P35 Nep KO mice, suggesting that the defect is established at early postnatal development and persists over the lifetime of the mouse (**Supplementary Figure S2**).

**Figure 5.**
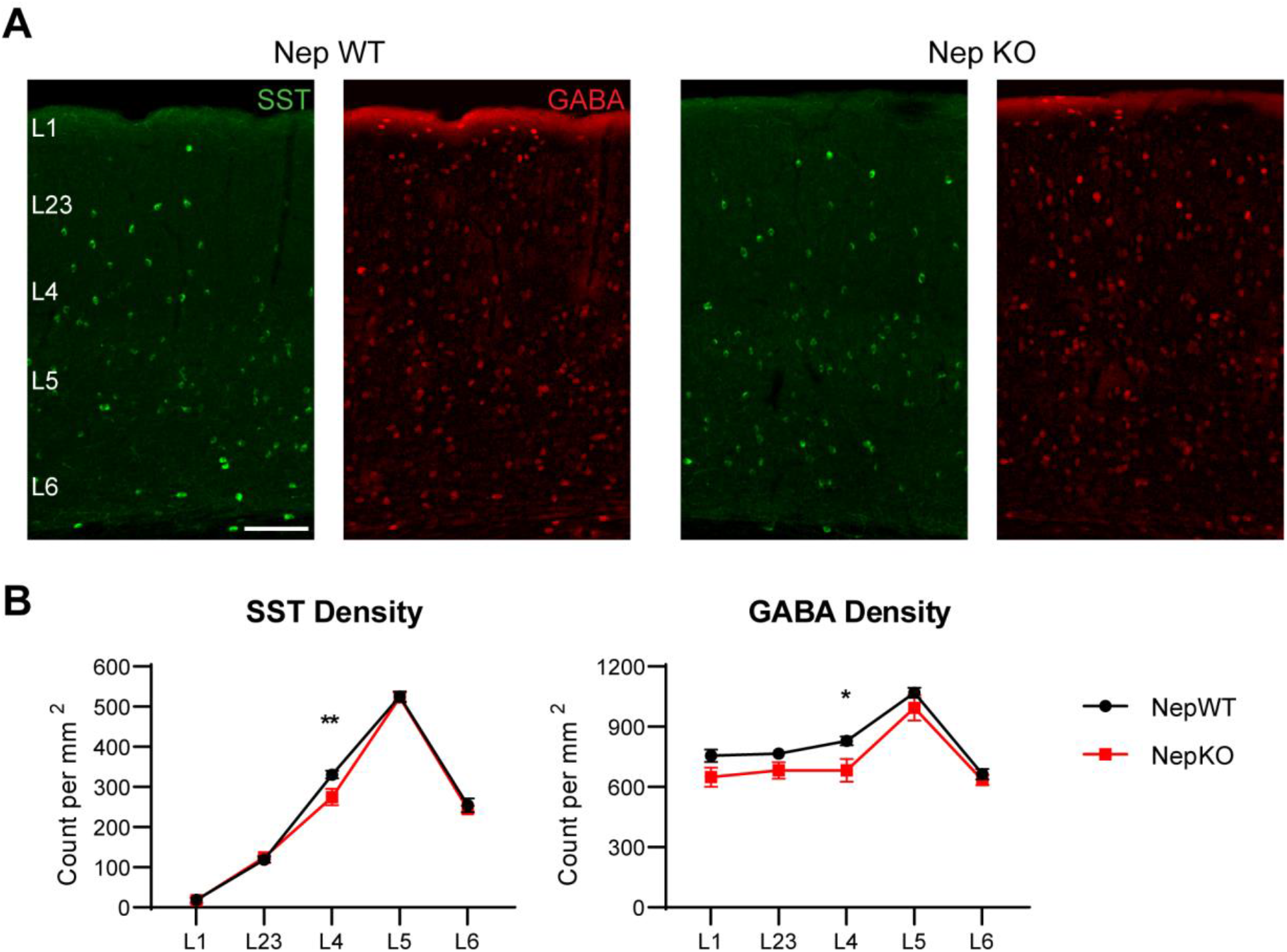
Reduced SST and GABA cells in L4 of Nep KO adult mice. **(A)** Representative images showing SST and GABA staining in the Nep WT and KO. (Scale bar, 100 μm) **(B)** Quantification of SST+ and GABA+ cells. There is a 17% reduction in the density of SST and GABA neurons in L4. (Nep WT n = 8, KO n = 6; Repeated Measures Two-way ANOVA with Bonferroni post-hoc test, * *p* < 0.05, ** *p* < 0.01)

### Nep KO mice show reduced activation of L5 and delayed maturation of optomotor acuity

L4 of the primary visual cortex is an important thalamorecipient layer, and the inhibitory neurons of L4, particularly PV neurons, exert strong feed-forward inhibition onto excitatory neurons, gating the spread of activity throughout the cortex (Cruikshank et al., 2007). The decrease in L4 inhibitory cells in Nep KO mice may reduce the strength of inhibitory gating and increase the spread of activity in the cortex. Voltage-sensitive dye imaging (VSDI) of acute coronal slices allows monitoring of the effect of macroscopic changes in the visual circuitry on activity.

Stimulation of the binocular portion of the primary visual cortex caused a rapid spread of depolarization from the white matter to the pia in a vertical column perpendicular to the pial surface (**Supplementary Figure S3A**). Then the signal traveled horizontally, with the greatest spread of activity in L23 which created an upper layer-heavy fan-shaped pattern. Responses peaked around 10 ms after stimulation (time, t = 110 ms) and fell back down to baseline levels over the course of a 511-ms recording (**Supplementary Figure S4B**). After imaging, the cortical slices were fixed in 4% PFA and stained with DAPI (**Figure 6A**), and this information was overlaid onto VSDI images, allowing more accurate delineation of cortical layers.

**Figure 6.**
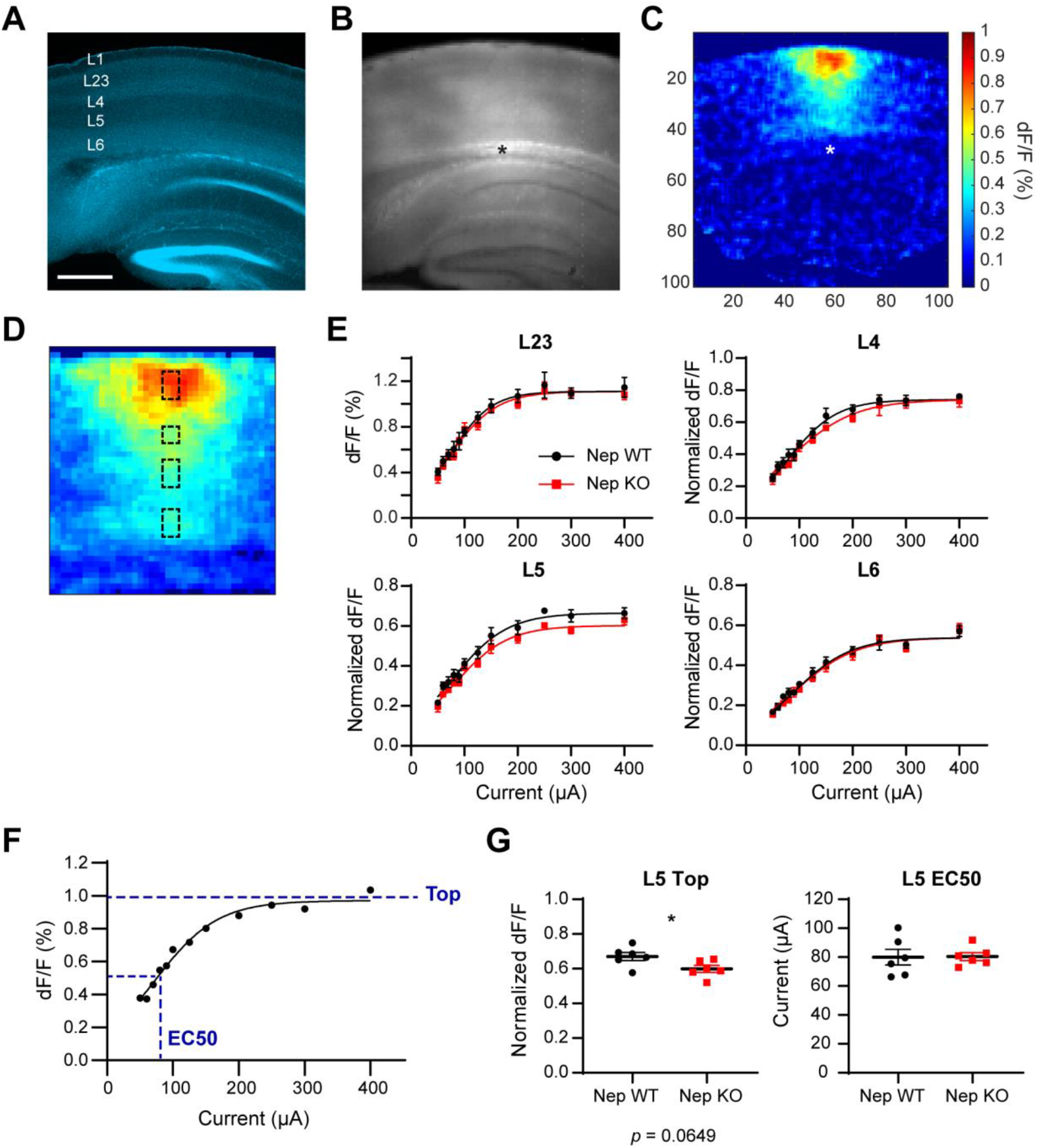
Voltage-sensitive dye imaging (VSDI) shows reduced activation of L5 in Nep KO mice. **(A)** DAPI staining of post-VSDI slices, which allows more accurate delineation of cortical layers. (Scale bar, 500 μm) **(B)** The background dye fluorescence of the same slice imaged during the VSDI experiment. The asterisk shows the site of activation. **(C)** dF/F heatmap of the slice 10 ms after white matter stimulation. **(D)** An example of flattened peak map at 100 μA stimulation. The four dashed boxes indicate ROIs that were averaged for the laminar analysis (L23, L4, L5, and L6 from the top). **(E)** Averaged current/response curves for the four laminar ROIs. (WT n = 6 from 3 mice, KO n = 6 from 3 mice) **(F)** An example current/response curve showing the peak dF/F after various stimulus intensities for a L23 ROI. A sigmoidal curve was fit through the data points, and two parameters were extracted from each fit, Top (the upper asymptote of the sigmoid function) and EC50 (the current at half maximal response, or 50% of Top). **(G)** Comparisons of parameters Top and EC50 of the laminar ROIs in Nep WT and KO. L23, L4, and L6 responses were not different in the WT and KO (**Supplementary Figure S4A**). Nep KO slices showed a 10% drop in L5 maximal response (0.67 ± 0.02 vs. 0.60 ± 0.02; t-test, * *p* < 0.05)

Using this information, we measured the peak responses of four regions of interest (ROIs) at the center of L23, L4, L5, and L6 over increasing stimulation intensities (**Figure 6D**). The average dF/F increased linearly with increasing stimulation current until the responses saturated around 150–200 μA (**Figure 6E**). The current/response curves could be fit with a sigmoidal curve with high goodness-of-fit (R^2^ > 0.9). Two parameters were extracted and compared for each layer: Top, the upper asymptote of the sigmoid function and the maximal response of the layer and EC50, the value of current at which half maximal response occurs (**Figure 6F**). Notably, L5 of Nep KO slice showed a 10% drop in the maximal response compared to WT (**Figure 6G**; 0.67 ± 0.02 and 0.60 ± 0.02, t-test, *p* = 0.0434). There were no differences in L23, L4, or L6 between the WT and KO (**Supplementary Figure S4**).

To assess where there might be any functional consequence of the reduced activity of L5 in Nep KO, we measured the development of optomotor acuity with optomotor task (Prusky et al., 2004). Over the course of early postnatal development, mice develop an innate ability to track moving visual stimuli with corresponding head motion. In this test, mice are placed on a platform surrounded by screens with slowly rotating sinusoidal gratings of various spatial frequencies (**Figure 7A**). From eye opening around P12–14, WT optomotor visual acuity rises steadily to adult acuity near the time of weaning (P21). Nep KO mice, however, did not reach adult acuity until P30, indicating that aspects of visual function and development are impaired by the deficiency of Nep protein (**Figure 7B**).

**Figure 7.**
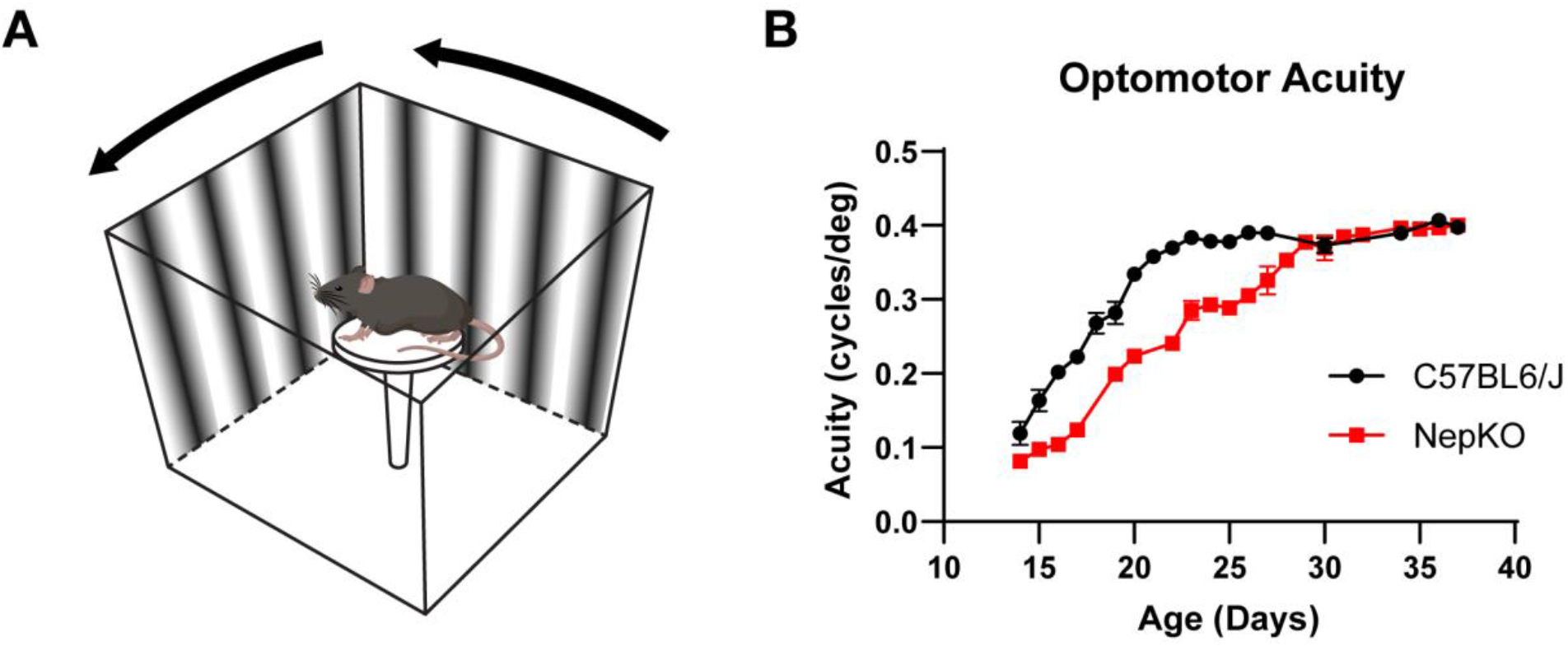
Nep KO mice show delayed maturation of optomotor acuity. **(A)** A schematic depicting the optomotor acuity test set-up. A mouse is placed in the center platform surrounded by computer monitors with moving sine gratings. **(B)** Optomotor acuity test of postnatally developing C57BL6/J and Nep KO mice. Nep KO mice showed a 10-day delay in visual acuity maturation. The number of mice used for each age data point ranged from n = 4 to 20 (**Supplementary Table S1**).

## 5 Discussion

In the current study, we investigated the role that Nep endopeptidase plays in the maturation of the mouse visual cortex. Nep is highly expressed in the young cortex at P10 and its expression is rapidly downregulated after eye-opening. L5 of the cortex expresses an especially high level of Nep, mostly in inhibitory neurons but also at low levels in many excitatory neurons. Potentially due to these laminar differences in the developmental expression, the deficiency of Nep leads to several layer-specific phenotypes: a decrease in the number of L4 inhibitory neurons and a reduction in the maximal activation of L5 after white matter stimulation. Contrary to our initial hypothesis that Nep may play a role in the development or modification of the PNNs, given their physical location on PV cell plasma membrane, we did not find any alterations in the ratio of PNN formation on PV neurons. Behaviorally, these changes manifested as a delay in the development of optomotor acuity.

### Nep as a potential molecular marker of PV neuron subtype in the cortex

Inhibitory neurons of the cortex vary widely in their morphology, connectivity, electrophysiological properties, and molecular markers (Markram et al., 2004; DeFelipe et al., 2013; Kepecs and Fishell, 2014). PV-expressing inhibitory neurons account for up to half of all inhibitory neurons in the cortex, yet the molecular classification is imperfect (Hu et al., 2014). There are at least two functional subtypes of PV-expressing neurons with distinct electrophysiological and morphological properties, axo-axonic chandelier cells and soma-targeting basket cells. Some studies have found PV subtypes that are born at different embryonic ages and differentially affected by learning (Donato et al., 2013, 2015). In addition, PV expression only begins around P20 and slowly ramps up (Río et al., 1994; Ye and Miao, 2012), making it unfit for early labeling or perturbation of the cells. Ideal novel markers for functional PV neuron subtypes would show higher specificity and earlier expression for reporter generation and genetic manipulation.

Rossier and colleagues showed that Nep is expressed by a subpopulation of PV neurons associated with PNNs and higher fast-spiking activity, suggesting that Nep could be a much-needed marker of a PV neuron subtype (Rossier et al., 2015). However, the developmental expression pattern of Nep in the visual cortex presented in the current study shows that Nep expression is not limited to inhibitory neurons at P10 and potentially at earlier ages (**Figure 3**). Such developmental narrowing of expression pattern is not unique to Nep; for example, previous work has shown that neurogranin and K_v_3.1, whose expressions are limited to excitatory cells and inhibitory neurons respectively in the mature cortex, are transiently expressed by more cell types in younger brains (Kalish et al., 2020).

This broader expression of Nep in the younger cortex prevents its use as an early reporter of a PV neuron subtype using the standard Cre-loxP strategy, but it may be possible with a modified approach. While the signal for Nep transcripts inside inhibitory neurons was highly concentrated and seemed to fill the entire cell body (**Figure 3C**; bottom blue arrows), excitatory neurons contained only a few speckles of puncta per cell, indicating low expression (bottom blue arrowheads). This stark difference in the level of Nep expression between excitatory neurons and the subset of inhibitory cells (which are likely to mature into PV-expressing neurons in the future) can be utilized to create a reporter system using mutant loxP sites with reduced recombination efficiency (Klinger et al., 2010; Liu and Sanes, 2017). Under this system, fluorophore expression can be limited to cells with the highest expression of Nep, allowing specific labeling of the PV neuron subtype that is of high interest to researchers studying critical period plasticity and PNNs (Beurdeley et al., 2012; Cabungcal et al., 2013; Sorg et al., 2016; Fawcett et al., 2019).

### Nep may clear Aβ accumulation from PV synapses in L5

The expression of Nep at P10 was highly laminar, with much higher expression in the lower layers, for both inhibitory and excitatory neurons (**Figure 3**). This potentially led to the laminar alterations in cortical response after white matter stimulation (**Figure 6G**). While the majority of Nep is produced in inhibitory neurons at P10 and in PV neurons at P40 (**Figure 3C**), in agreement with previous work (Rossier et al., 2015; Tasic et al., 2018), other groups have shown that Nep protein localizes primarily to the presynaptic terminals in the mouse hippocampus (Fukami et al., 2002; Iwata et al., 2004) and surrounding the cell bodies of L5 pyramidal neurons in the somatosensory cortex (Pacheco-Quinto et al., 2016). PV neurons are characterized by their perisomatic boutons that a create dense net of innervation around the cell bodies of target cells (Freund and Katona, 2007), and PV-immunopositive fibers (**Figure 4A & Supplementary Figure S2A**) and Syt2-positive PV boutons (Sommeijer et al., 2012) are highly concentrated in L5 of the cortex. Together, these data suggest that Nep produced by PV neurons are transported to the presynaptic terminals apposing pyramidal neuron cell bodies in L5 (**Figure 1B**; right image), potentially modulating the function of the inhibitory synapses by regulating the level of Aβ.

Aβ is the most prominent and well-researched substrate of Nep in the brain and has been shown to be released from synapses into the extracellular space in an activity-dependent manner (Kamenetz et al., 2003; Cirrito et al., 2005). At the synapse, Aβ plays a dose-dependent hormetic role in synaptic plasticity. At pathological nanomolar concentrations, Aβ impairs synaptic function and inhibits hippocampal long-term potentiation (LTP) (Walsh et al., 2002). However, at physiological high picomolar concentrations (Cirrito et al., 2003), Aβ has been shown to enhance LTP in mouse hippocampal neurons, and correspondingly improve their reference and contextual memory (Puzzo et al., 2008, 2011). Notably, application of Nep inhibitor thiorphan to rat hippocampal and cortical neuron cultures elevated the level of secreted Aβ in the media, and increased activity-driven synaptic vesicle cycling at the synapses (Abramov et al., 2009; Lazarevic et al., 2017). These studies show that acute inhibition of Aβ catabolism by Nep can enhance synaptic transmission by increasing the release probability.

We found that Aβ accumulates in the visual cortex in the absence of Nep, even from a juvenile age (**Figure 2B**). Thus, increased Aβ levels at the synapses of PV neurons in Nep KO mice could lead to the reduction of L5 cortical response. The accumulation of soluble Aβ at the PV synapses in L5 may lead to increased release of GABA onto the pyramidal cell bodies, dampening L5 response after cortical stimulation. This potential change in PV synaptic activity may underlie the impairment of ocular dominance plasticity in APPswe/PS1 mice, a mouse model for Alzheimer’s disease (William et al., 2012). Expression limited to PV neurons means that deficiency in Nep can alter PV synapses specifically, with implications in excitatory/inhibitory (E/I) balance important for cortical plasticity (Hensch, 2005). It is also interesting that fast-spiking PV neurons which are more vulnerable to activity-dependent Aβ accumulation are the primary producers of Nep, suggesting a potential negative feedback mechanism regulating the level of Aβ and optimal synaptic activity in the cortex.

These data suggest that the deficiency of Nep at L5 during early postnatal development in Nep KO mice may lead to impairments in L5 activation. The relatively small size of reduction in response (∼10% decrease; **Figure 6G**) could be due to the age of the mice at the time of VSDI experiments. Mice were tested during the time window P25–30, when the developmental ramping down of Nep expression has progressed substantially (**Supplementary Figure S1**), which may have masked the difference between Nep WT and KO. Because the expression of Nep is highest at P10 or younger, cortical response at younger ages should be further investigated.

### Developmental cortical feedback for optomotor task in Nep KO mice may be impaired

Important functional consequence of Nep KO is the delay in optomotor acuity development. While WT mice reach maximum optomotor acuity around P25, Nep KO only reached this level at P35, showing a 10-day delay (**Figure 7**). Both optomotor response, where the head movement tracks the slow horizontally moving stimuli, and a related phenomenon called optokinetic response, in which the eye movements track the stimuli, are mediated by subcortical visual pathways from the retinal ganglion cells to the accessory optic system (AOS) in the midbrain, particularly a structure called NOT-DTN (nucleus of the optic tract-dorsal terminal nucleus) (Giolli et al., 2006). While an intact visual cortex is not necessary for the optomotor task, as bilateral lesion or pharmacological inactivation does not affect the response in adult mice (Douglas et al., 2005), cortical inputs are necessary for the plasticity. For example, a short monocular deprivation increases the optomotor response of the non-deprived eye in adult mice, but this experience-dependent plasticity is abolished when the visual cortex is silenced (Prusky et al., 2006). Likewise, vestibular lesion-dependent potentiation of optokinetic response requires the activation of the visual cortex (Liu et al., 2016). The study showed that a subpopulation of L5 pyramidal neurons makes functional projections to NOT-DTN and mediates the change in response.

Given the importance of L5 cortical inputs for optomotor plasticity, it is possible that the deficiency of Nep in L5 of the visual cortex in Nep KO mice and the weakening of L5 activation may alter its feedback onto AOS, leading to changes in developmental trajectory of the optomotor response. The discrepancy in the optomotor acuity of WT and Nep KO mice begins after eye opening and diminishes as the mice approach P35 (**Figure 7B**), matching the timeline of Nep downregulation in the visual cortex (**Figure 3**). The role of cortical input during the maturation of optomotor acuity in mice has not been investigated, and our current results support such a role.

### Decrease in the number of inhibitory neurons in the thalamorecipient layer of the cortex

Nep KO mice showed a 17% reduction in the number of inhibitory neurons specifically in L4, the principal thalamorecipient layer of the cortex (**Figures 4 and 5**). This phenotype was seen as early as P17 and also in P35 mice, suggesting that it is established early in life and continues into adulthood (**Supplementary Figure S2**). In many other mouse models of disrupted laminar organization, inhibitory neurons are mislocalized to aberrant layers as a result of impairments in migration or disruptions in cell-fate specification during neurogenesis (Liodis et al., 2007; Vogt et al., 2014; Bartolini et al., 2017). Unlike their excitatory counterparts, cortical interneurons are born in the ventricular zones of ganglionic eminences of the embryonic brain and migrate tangentially in distinct migratory streams before migrating radially into the cortical plate (Pla et al., 2006; Lopez-Bendito et al., 2008; Faux et al., 2012), creating a window of high vulnerability for defective migration. Nep was shown to regulate the migration of prostatic cancer cells via inhibiting the phosphorylation of focal adhesion kinase (Sumitomo et al., 2000, 2004), and it is possible that Nep on migrating inhibitory neurons in the embryonic brain may guide their migration and laminar distribution. Currently it is not known whether Nep is expressed in the embryonic brain, but the downward trajectory of its developmental expression in the visual cortex, as well as the broad early expression pattern, (**Figures 2A and 3**) supports this hypothesis.

Given the important role of L4 in receiving and gating inputs from the thalamus, we hypothesized that the spread of activation throughout the cortex might be altered in the Nep KO due to the reduction in the number of inhibitory neurons. However, we found no observable difference in the vertical or horizontal spread of depolarization from L4 after white matter stimulation (**Figure 6 and Supplementary Figure S4**). After the delivery of current pulse, activity spreads vertically from white matter to the pial surface, passing through L4, before spreading horizontally in a fan shape (**Supplementary Figure S3A**). There were no differences in the maximum activation of L23 or the horizontal width of activation in L23 and L4, indicating that the macroscopic circuitry of the visual cortex is not altered in the Nep KO. The peak dF/F of L5 occurs before the peak of L4 (**Supplementary Figure S3A**), and the 10% drop in its maximal activation (**Figure 6G**) cannot be construed as a consequence of reduced spread from L4 in this experimental set-up.

White matter stimulations used in the current study activate not only thalamocortical fibers but also other axons innervating the particular patch of the cortex, such as callosal fibers conveying input from the opposite hemisphere and other cortico-cortical connections, as well as potential antidromic activation of the cortex by axon collaterals of efferent axons. A more sensitive and more appropriate experimental condition to detect the functional consequences of decreased L4 inhibition in Nep KO mice would be a thalamocortical slice. A thalamocortical slice preserves the inputs from lateral geniculate nucleus (LGN) and allows specific activation of neurons receiving direct thalamic inputs such as PV neurons of L4 (MacLean et al., 2006). Future work probing the cortical circuitry of Nep KO mice can use thalamocortical stimulations to overcome the limitations of the current study.

## 7 Supplementary Figure Legend

**Supplementary Figure S1.**
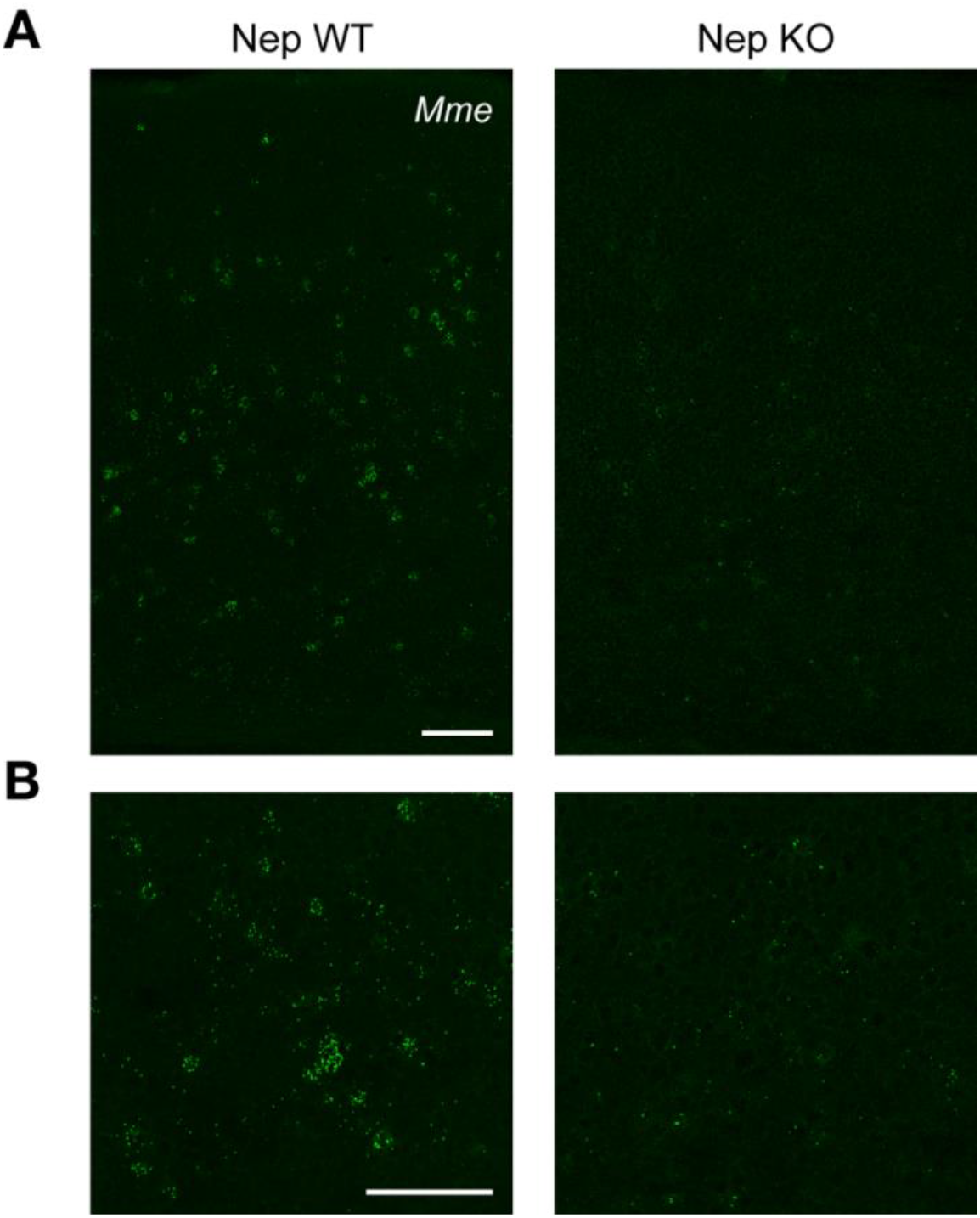
*Mme* probe signal is absent in Nep KO mouse visual cortex. (**A**) Representative images showing *Mme in situ* hybridization signals in P25 Nep WT and Nep KO visual cortex. Punctate signals are clearly visible inside select cells throughout the cortical layers in Nep WT while Nep KO only shows sparse background staining. (Scale bar, 100 μm) (**B**) Zoomed-in versions of the above images for easier inspection of the *Mme* puncta. (Scale bar, 100 μm)

**Supplementary Figure S2.**
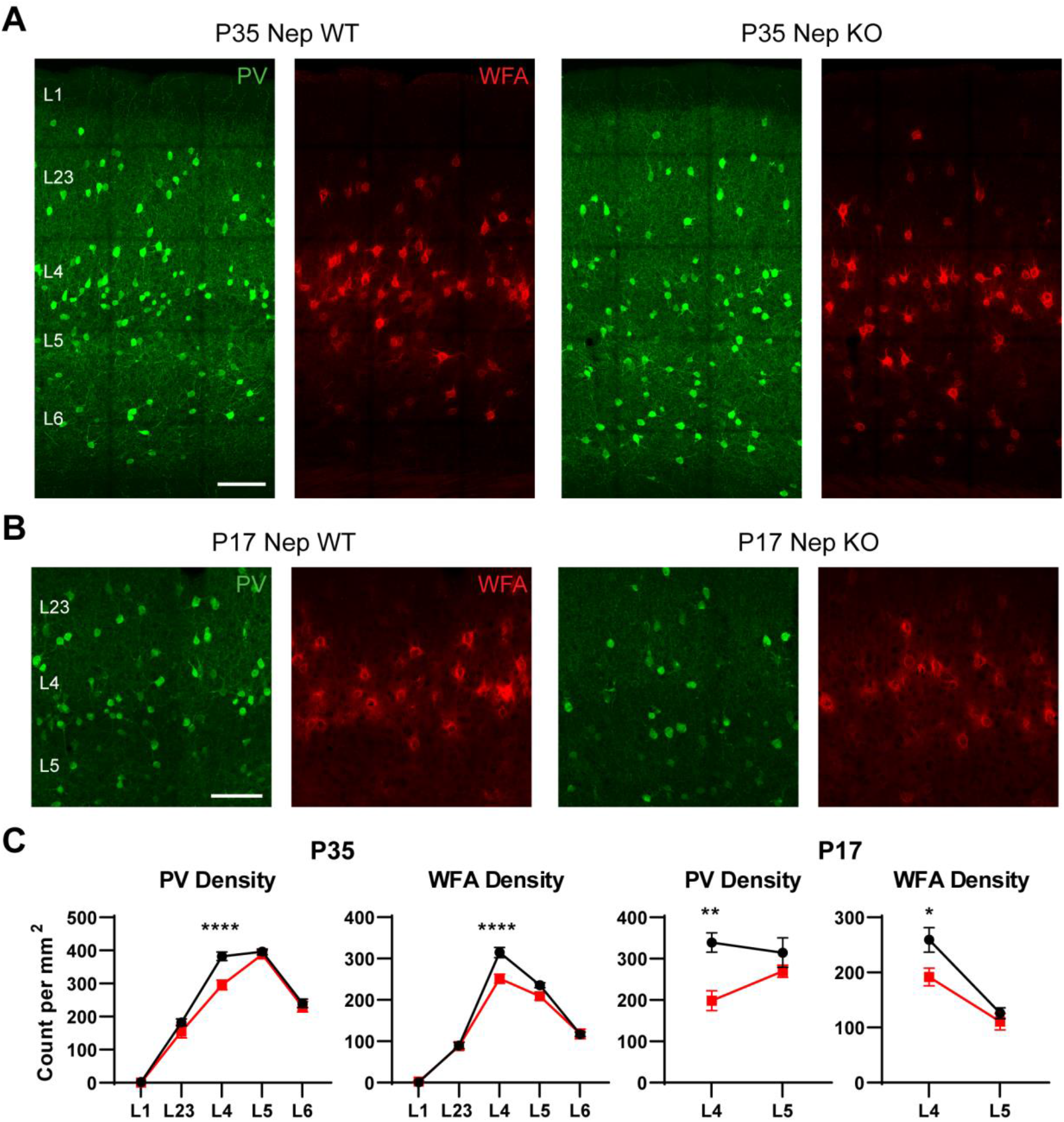
L4 PV neurons and WFA cells are decreased in P35 and P17 Nep KO mice. (**A**) Representative images of P35 Nep WT and KO sections with PV and WFA immunostaining. (Scale bar, 100 μm) (**B**) Representative P17 images centered at L4. (Scale bar, 100 μm) (**C**) Quantification of PV and WFA densities. Both P35 and P17 Nep KO mice show decreased PV and WFA cells in L4 of the visual cortex. (P35 WT n = 8, KO n = 5; P17 WT n = 4, KO n = 4; Repeated Measures Two-way ANOVA with Bonferroni post-hoc test, * *p* < 0.05, ** *p* < 0.01, **** *p* < 0.0001)

**Supplementary Figure S3.**
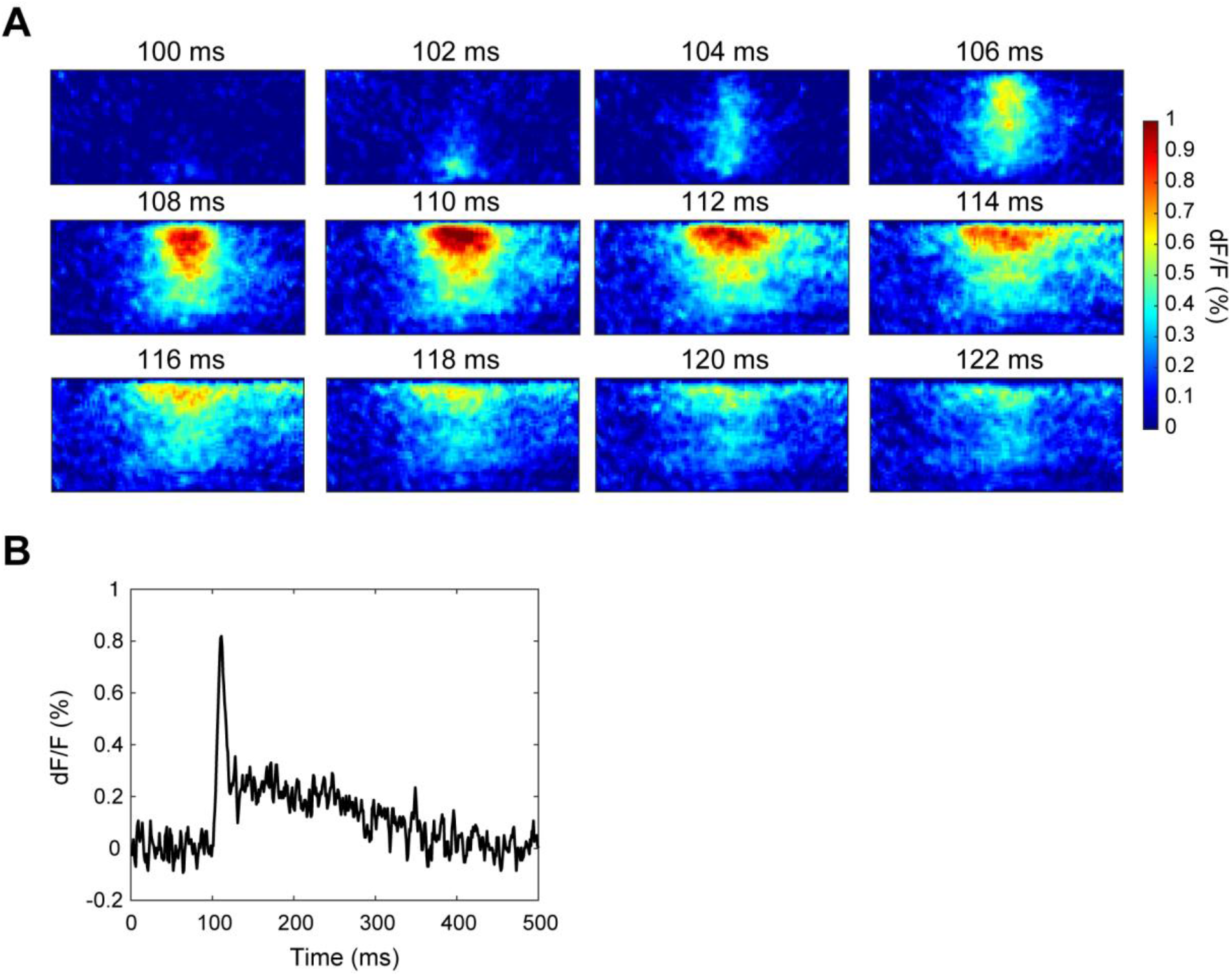
Activation pattern of acute visual cortical slices after white matter stimulation. (**A**) dF/F heatmap showing the vertical and horizontal spread of depolarization through the cortex during the first 20 ms of stimulation. The images were flattened to allow for laminar analysis. (**B**) The dF/F response at the center of L23 over time. The white matter stimulation occurs at 100 ms.

**Supplementary Figure S4.**
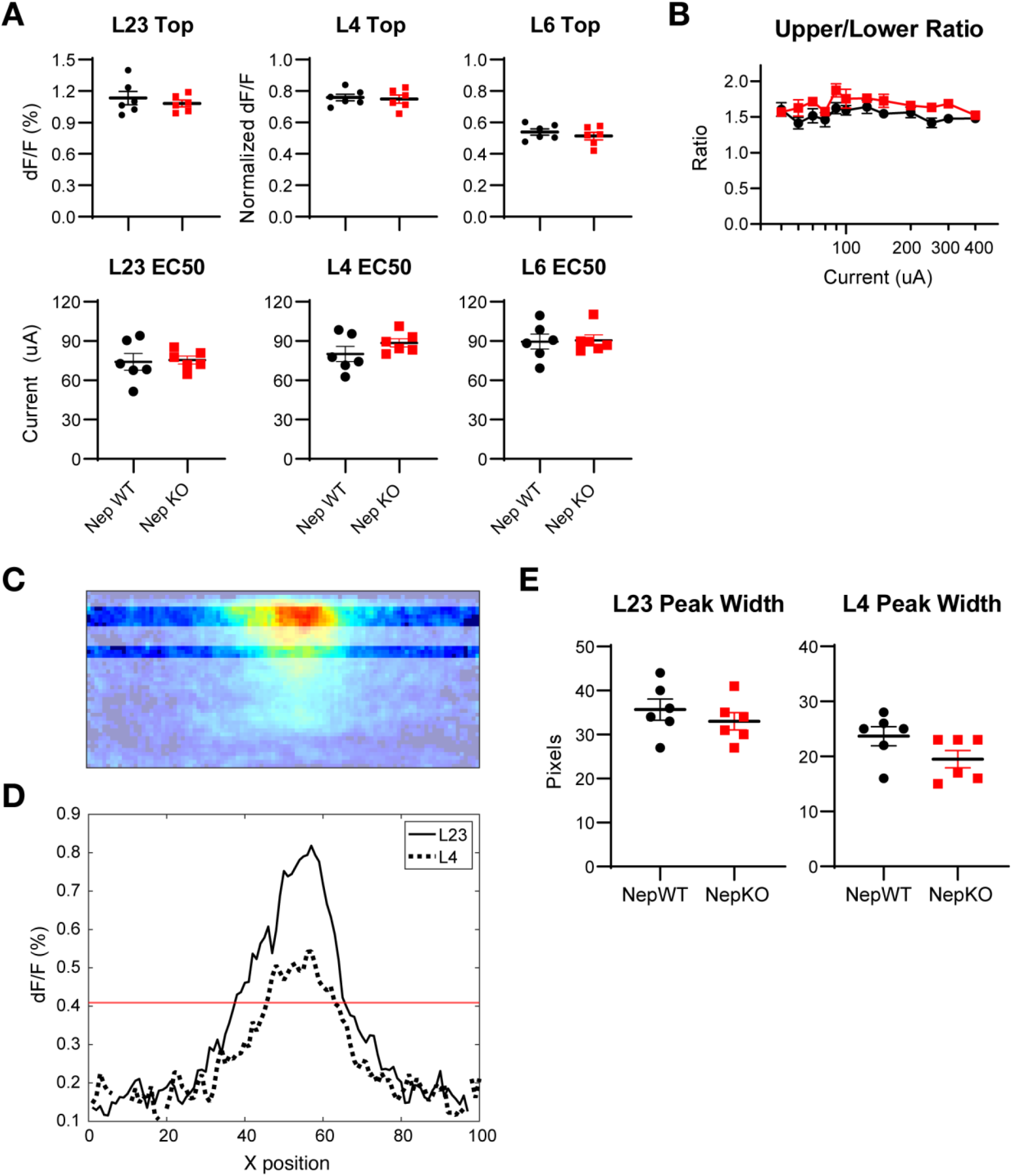
L23, L4, and L6 vertical spread and horizontal spread are not altered in Nep KO visual cortex. (**A**) Comparisons of parameters Top and EC50 of the laminar ROIs in Nep WT and KO. L23, L4, and L6 responses were not different in the WT and KO. (**B**) The ratio of L23 and L5 peak responses as a function of white matter stimulus intensity. All results are expressed as mean ± SEM (WT n = 6 from 3 mice, KO n = 6 from 3 mice). Previously, a mutant mouse model with enhanced inhibition (MeCP2 KO) exhibited decreased spread of activity to the upper layers after white matter stimulations, which led to a decreased ratio of upper- and lower-layer activity (Durand et al., 2012). To see whether there might be subtle differences in lower to upper layer spread that were drowned out by individual variability, we calculated the upper/lower layer ratio at each stimulus intensity. There were no interactions between the ratio and current as seen in Durand et al. and no difference between WT and KO at any current intensity. (**C–E**) The horizontal spread is not altered in Nep KO visual cortex after white matter stimulation (**C**) An example peak map showing the regions analyzed for the horizontal spread analysis. The two bands with higher opacity are L23 and L4. (**D**) An example plot showing the averaged values of L23 and L4 at each X position across all the columns of the above image. The dF/F signal of each X-position in the image was averaged. The red line indicates the threshold of half maximum in L23. (**E**) The width of the spread was defined as the number of continuous pixels higher than the threshold. There were no differences in the width of L23 horizontal spread and a small decrease in the width of L4 horizontal spread in the KO, but this was not statistically significant (*p* = 0.1058).

**Supplementary Table S1.** Number of animals used in optomotor task.

| Age | C57BL6/J | Nep KO | Age | C57BL6/J | Nep KO | Age | C57BL6/J | Nep KO |
| --- | --- | --- | --- | --- | --- | --- | --- | --- |
| 14 | 11 | 4 | 22 | 17 | 18 | 30 | 4 | 10 |
| 15 | 10 | 14 | 23 | 6 | 10 | 31 | 0 | 7 |
| 16 | 19 | 4 | 24 | 20 | 4 | 32 | 0 | 14 |
| 17 | 6 | 17 | 25 | 5 | 7 | 34 | 7 | 11 |
| 18 | 7 | 0 | 26 | 4 | 11 | 35 | 0 | 7 |
| 19 | 7 | 4 | 27 | 15 | 10 | 36 | 9 | 7 |
| 20 | 14 | 17 | 28 | 0 | 7 | 37 | 5 | 7 |
| 21 | 10 | 0 | 29 | 0 | 7 |  |  |  |

## 8 Author Contributions

HB and TH designed the study; HB performed the experiments and data analyses; HB drafted the manuscript, and HB and TH revised the manuscript.

## Acknowledgments

We thank N. De Souza, R. Woo, and K. Hoffmann for animal care and maintenance; G. Gunner from Neurobehavioral Core for performing optomotor task experiments; Harvard Center for Biological Imaging (HCBI) and CBS imaging core for confocal imaging.

